# Comparative pathogenicity of SARS-CoV-2 Omicron subvariants including BA.1, BA.2, and BA.5

**DOI:** 10.1101/2022.08.05.502758

**Authors:** Tomokazu Tamura, Daichi Yamasoba, Yoshitaka Oda, Jumpei Ito, Tomoko Kamasaki, Naganori Nao, Rina Hashimoto, Yoichiro Fujioka, Rigel Suzuki, Lei Wang, Hayato Ito, Izumi Kimura, Isao Yokota, Mai Kishimoto, Masumi Tsuda, Hirofumi Sawa, Kumiko Yoshimatsu, Yusuke Ohba, Yuki Yamamoto, Tetsuharu Nagamoto, Jun Kanamune, The Genotype to Phenotype Japan (G2P-Japan) Consortium, Keita Matsuno, Kazuo Takayama, Shinya Tanaka, Kei Sato, Takasuke Fukuhara

## Abstract

Unremitting emergence of severe acute respiratory syndrome coronavirus 2 (SARS-CoV-2) variants imposes us to continuous control measurement. Given the rapid spread, new Omicron subvariant named BA.5 is urgently required for characterization. Here we analyzed BA.5 with the other Omicron variants BA.1, BA.2, and ancestral B.1.1 comprehensively. Although *in vitro* growth kinetics of BA.5 is comparable among the Omicron subvariants, BA.5 become much more fusogenic than BA.1 and BA.2. The airway-on-a-chip analysis showed that the ability of BA.5 to disrupt the respiratory epithelial and endothelial barriers is enhanced among Omicron subvariants. Furthermore, in our hamster model, *in vivo* replication of BA.5 is comparable with that of the other Omicrons and less than that of the ancestral B.1.1. Importantly, inflammatory response against BA.5 is strong compared with BA.1 and BA.2. Our data suggest that BA.5 is still low pathogenic compared to ancestral strain but evolved to induce enhanced inflammation when compared to prior Omicron subvariants.

## Main

In recent months, multiple omicron sub-lineages of severe acute respiratory syndrome coronavirus 2 (SARS-CoV-2) have emerged^1^ and impose continually public concerns for COVID-19 control measurement. The subvariants named BA.4/BA.5 are firstly isolated in South Africa^2^. Now they have been detected in dozens of countries worldwide, with a combined frequency of over 50% in recent weeks at the time of writing the initial manuscript of this study in July 2022. As of August 2022, BA.5 has outcompeted the original BA.2 and becomes dominant variant in the world. The raising cases of COVID-19 by BA.4/BA.5 indicate that they acquire the enhanced transmission ability compared to the sister linages, BA.1 and BA.2. Indeed, as shown in a recent report from Portugal where outbreak by BA.5 was occurred^3^, morbidity of BA.5 is higher than that of BA.1 variants. To answer potential urgency for COVID-19 wave, the several groups conducted the ability of immunoprophylaxis conferred by vaccination or infection with previous variants. The recent reports showed that BA.4/BA.5 substantially escape from neutralizing antibodies induced by vaccination or infection^4–8^. By genome sequencing and evolutionary analyses, BA.4 and BA.5 are more similar to BA.2 than to the BA.1 strain that surged in late last year. We have been shown that viral spike protein is one of the major virulence determinants^9–12^.

BA.4 and BA.5 have identical spike protein and carry their own unique mutations, including L452R that exhibited the enhanced fusogenic activity and pose resistant to the immunity induced by the infection with early variants^13^. This observation is consistent with the series of our studies using the recombinant viruses replacing the spike protein gene from the ancestral early pandemic variants. However, virological characters of BA.5 strain isolated from COVID-19 patients has not yet fully defined. Here, we employed the indicated omicron subvariants (BA.1 lineage, strain TY38-873, GISAID ID: EPI_ISL_7418017; BA.2 lineage, strain TY40-385, GISAID ID: EPI_ISL_9595859; BA.5 lineage, strain TKYS14631, GISAID ID: EPI_ISL_12812500)^14^ for investigating virological characters *in vitro* and *in vivo* model.

## Virological features of Omicron subvariants *in vitro*

To elucidate the virological characteristics of Omicron subvariants, we obtained BA.1 isolate (strain TY38-873), BA.2 isolate (strain TY40-385), and BA.5 isolate (strain TKYS14631). A D614G-bearing early pandemic B.1.1 isolate (strain TKYE610670)^10^ was used as a control. Then we characterized their *in vitro* growth kinetics using the cell lines VeroE6/TMPRSS2^15^, Calu-3, and iPS cell-derived alveolar epithelial cells (**Fig. 1A, B**). In the cell lines VeroE6/TMPRSS2 and Calu-3, BA.2 and BA.5 subvariants are comparable replication to the B.1.1 but BA.1 showed low replication rate. In the iPS cell-derived alveolar epithelium cells, B.1.1, BA.1, and BA.5 exhibited the slightly high replication efficiency compared with the strain BA.2, suggesting that the replication property of Omicron subvariant BA.5 *in vitro* is similar to that of B.1.1. Although the growth kinetics of Omicron subvariants and B.1.1 in VeroE6/TMPRSS2 cells were comparable (**Fig. 1A**), the morphology of infected cells was quite different; B.1.1 formed larger syncytia than the Omicron subvariants (**Fig. 1C**). To further investigate syncytia formation of the Omicron subvariants, we generated the VeroE6/TMPRSS2 cells expressing either EGFP or mCherry and an equal amount of them were seeded for syncytia formation as a merged image (**Fig. 1D**). Within Omicron subvariants, the ability of syncytia formation of BA.1 is lower than that of BA.2 and BA.5. Consistent with observation of syncytia, efficacy of cleavage of S protein by B.1.1 infection was highest of all (**Fig. 1E**). Among the Omicron subvariants, BA.5 exhibited most efficient activity of S cleavage. These data suggest that even though Omicron subvariants are still less fusogenic than B.1.1 isolate, the subvariants are evolving toward efficient fusogenicity in VeroE6/TMPRSS2 cells. A previous study has shown that some ultrastructural changes are detected in SARS-CoV-2-infected cells at early phase of post infection^16^. One of the alterations is the presence of annulate lamellae (AL) in the cell sections. AL are characterized as stacks of ER-derived membranes highly arranged in parallel and containing nuclear pore complexes^17^. To investigate whether the difference in the extent of AL formation upon infection with the variants, we performed electron microscopic analysis (**Fig. 1F**) of SARS-CoV-2-infected infected VeroE6/TMPRSS2 cells at early phase of post infection (multiplicity of infection, m.o.i. = 0.01, 16 h.p.i). In uninfected cells, AL were not seen even at late phase (32 h.p.i) (**Fig. 1F**: mock). In contrast, by virus infection, several membranous structures (~ø80 nm) were gathered within electron-dense areas near the nucleus (**Fig. 1F**: B.1.1, BA. 1, BA.2) or at the expansion of the nuclear envelope (**Fig. 1F**: BA.5). They exhibited ultrastructural resemblance to cross sections of AL as shown in the earlier report^16^, suggesting that B.1.1 and Omicron subvariants are similar replication rate at the earlier stage of virus life cycle. Next, to evaluate the influence of viral infection on the respiratory epithelial and endothelial barriers, airway-on-a-chip was utilized; by examining the amount of virus that migrates from the airway channel to the blood vessel channel, the ability to disrupt the respiratory epithelial and endothelial barriers can be evaluated. In the Omicron subvariants, the largest amount of virus was detected in the blood vessel channel of BA.5-infected airway-on-a-chip. In addition, the B.1.1- and BA.5-infected airway-on-chip exhibited more severe disruption than the BA.1- and BA.2-infected ones, suggesting that BA.5 possesses a substantial barrier disruption capacity (**Fig. 1G**).

**Fig. 1.**
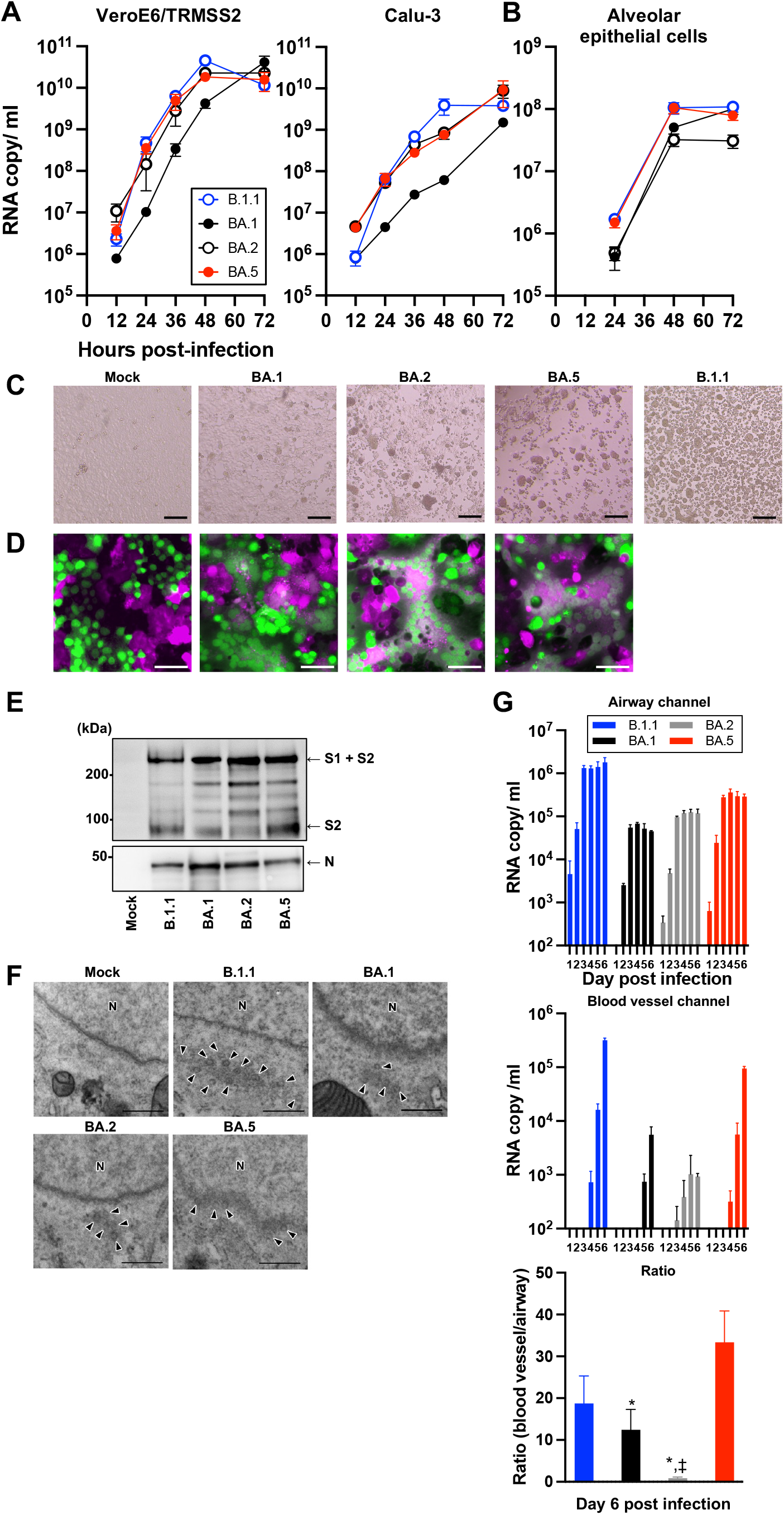
Virological features of Omicron subvariants *in vitro.* **A,B**, Growth kinetics of Omicron subvariants. The four clinical isolates of B.1.1, Omicron BA.1, BA.2, and BA.5 were inoculated into VeroE6/TMPRSS2, Calu-3 (**A**) and human alveolar epithelial (**B**) cells, and the copy number of the viral RNA in the supernatant was quantified by RT–qPCR. **C**, Bright-field images of infected VeroE6/TMPRSS2 cells (m.o.i. = 0.01) at 48 h.p.i. SARS-CoV-2 induced syncytia formation. EGFP- and mCherry-expressing VeroE6/TMPRSS2 cells were co-cultured at a 1:1 ratio and infected with B.1.1, BA.1, BA.2, and BA.5 isolates. Scale bars, 200 μm **D.** Syncytia formation was monitored by immunofluorescent microscopy at 48 h.p.i. Scale bars, 100 μm. **E,** Expressions of S protein (S1+S2 and S2) were examined by immunoblotting by monoclonal antibody against S upon infection with the respective SARS-CoV-2 variants in the VeroE6/TMPRSS2 cells. **F**, Ultrastructure of annulate lamellae (AL) in SARS-CoV-2 infected cells at 16 h.p.i. Arrowhead represents the membranous lamella which constitutes AL. Scale bar, 0.5 μm. **F**, The efficiency of S cleavage is evaluated by immunoblotting analysis. **G**, Airway-on-a-chip analysis. Medium containing SARS-CoV-2 was injected into the airway channel, which was then cultured for 6 days. Viral RNA in the supernatant of both the airway and blood vessel channels were quantified by RT–qPCR. Data are the average ± s.d. The ratio of viral invasion toward blood vessel channels were calculated (blood vessel channel/airway channel) on day 6 p.i. as shown percentages. (**A,B,G**). Assays were performed independently in two-time (**A**) or triplicate (**B,G**). In **G**, statistically significant differences between B1.1 and other variants (‡, *P* < 0.05), and between BA.5 and other variants (*, *P* < 0.05) were determined by two-sided Student’s *t* tests.

## Virological features of Omicron subvariants *in vivo*

To investigate the dynamics of viral replication *in vivo* and pathogenicity of Omicron subvariants, we utilized the established animal model using hamsters ^9–11^. Consistent with our previous study, B. 1.1-infected hamsters exhibited decreased body weight from day 2 post-infection (p.i.) (**Fig. 2A**). The change of the body weight of Omicron subvariants-infected hamsters was moderate compared with that of B. 1.1-infected hamsters and similar to the uninfected group. Importantly, the dynamics of weight changes of BA.5-infected hamsters was significantly different from that of the BA.2-infected and uninfected hamsters (**Fig. 2A**). We then quantitatively analyzed the pulmonary function of infected hamsters as reflected by three parameters, namely, enhanced pause (Penh) and the ratio of time to peak expiratory follow relative to the total expiratory time (Rpef), which are surrogate markers for bronchoconstriction or airway obstruction, and subcutaneous oxygen saturation (SpO_2_). As shown in **Fig. 2B–D**, the B.1.1-infected hamsters exhibited respiratory disorders according to these three parameters. In contrast, in BA.1-, BA.2- and BA.5-infected hamsters, the Penh value was significantly lower than those in B.1.1-infected hamsters (**Fig. 2B**), and the Rpef value was significantly higher than those in B.1.1-infected hamsters (**Fig. 2C**). As per SpO_2_ values, B.1.1- and BA.5-infected hamsters exhibited lower tendency than those of BA.1- and BA.2-infected hamsters (**Fig. 2D**). *In vivo* viral dynamics was analyzed by collecting the oral swab of infected hamsters at indicated timepoints (**Fig. 2E**). On day 1 p.i., the viral loads of Omicron subvariants-infected hamsters were significantly lower than those of B.1.1-infected hamsters. These data suggest that Omicron subvariants are less pathogenic than the B.1.1, but BA.5 is evolving toward slightly increased pathogenicity.

**Fig. 2.**
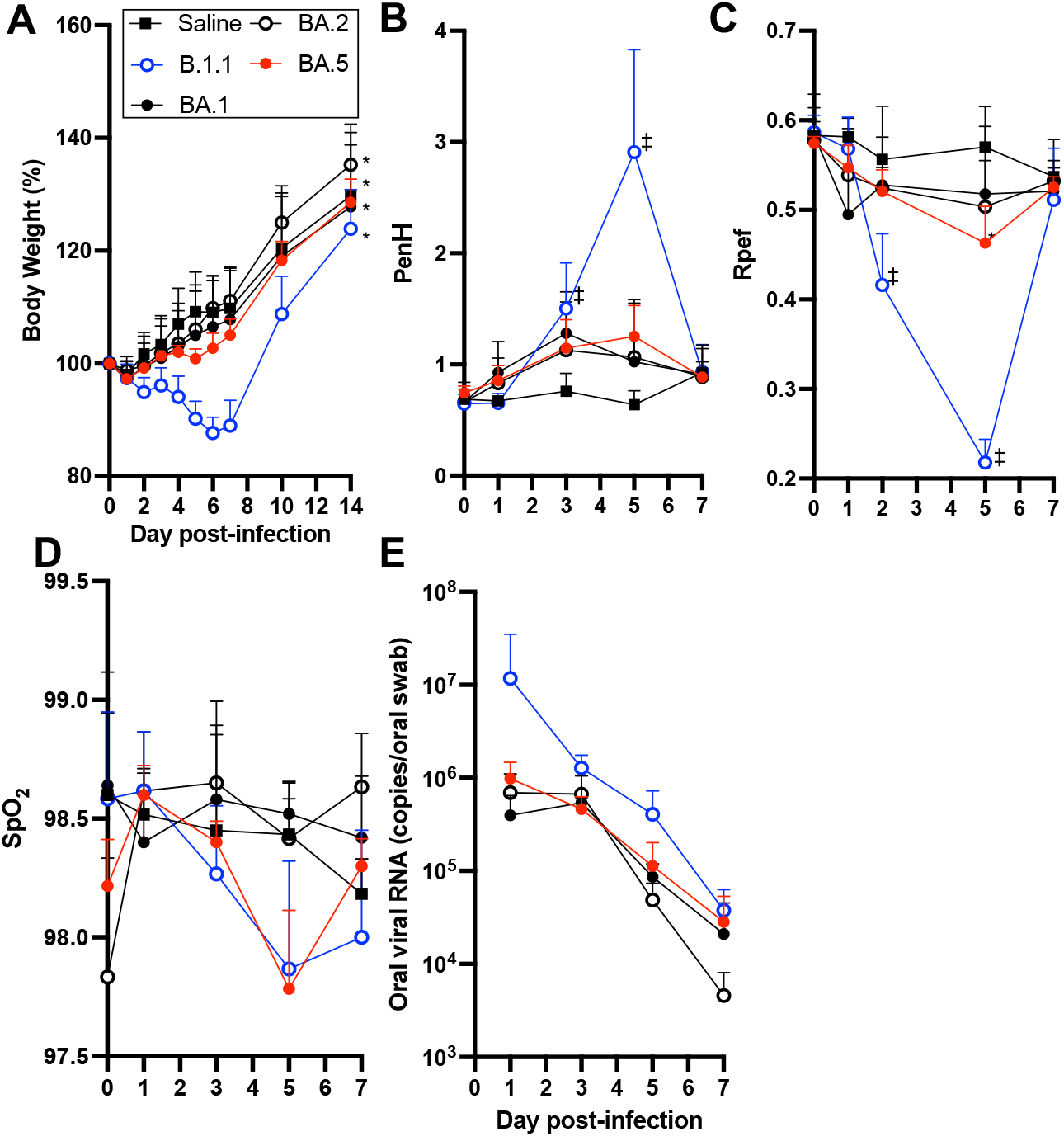
Time-course dynamics of Omicron subvariants *in vivo*. Syrian hamsters were intranasally inoculated with saline (*n* = 6, uninfected control), B.1.1 (*n* = 6), BA.1 (*n* = 6), BA.2 (*n* = 6) and BA.5 (*n* = 6). Body weight (**A**), Penh (**B**), Rpef (**C**), SpO_2_ (**D**), and viral RNA load in the oral swab (**E**) were routinely measured as indicated in the graph. Data are the average ± s.e.m. In **A**, statistically significant differences between BA.5 and other variants or saline across timepoints from on day 1 p.i. to on day 7 p.i. were determined by multiple regression (*, *P* < 0.05). The family-wise error rates calculated using the Holm method are indicated in the figure. In **B**-**D**, statically differences B.1.1 and other variants or saline were tested with Tukey’s multiplicity correction (‡, *P* < 0.05). Statically differences BA.5 and other variants or saline were tested with Tukey’s multiplicity correction (*, *P* < 0.05).

We next assessed viral spreading in the respiratory tissues and, thus, collected the trachea and the lung on day 2 and 5 p.i. (**Fig. 3A**). In the upper trachea of infected hamsters, epithelial cells were sporadically positive for viral N protein on day 2 p.i., but there were no significant differences among B.1.1 and Omicron subvariants (**Fig. S1B**). As per lung, we investigated the viral spreading in the separate region, hilum and periphery. In the lung hilum, although viral RNA copies (**Fig. 3A**, left) in the lung hilum of BA.1 were approximately 10-fold lower than those of B.1.1, BA.2, and BA.5 on day 2 p.i., viral RNAs of all omicron subvariants were significantly lower than those of B.1.1 on day 5 p.i. In contrast to the hilum, viral RNA copies (**Fig. 3A**, middle) and titers (**Fig. 3A**, right) in the periphery was slightly different among Omicron subvariants. Large amounts of viral load were detected from BA.5 on day 2 p.i. that is comparable to B.1.1. To further characterize virus spread by Omicron subvariants, immunohistochemical (IHC) analysis of viral N protein was conducted using the specimens of respiratory system. On day 2 p.i., the N protein was observed in the alveolar space around the bronchi/bronchioles in the B.1.1-infected hamsters (**Fig. 3B**, top panel). In Omicron BA.2- and BA.5-infected hamsters, the N protein was observed in the alveolar space with less extent than those of B.1.1. The N proteins strongly remain in lobar bronchi in the BA.5-, but not BA.2-, infected hamsters (**Fig. 3B**, BA.5: bottom panel, BA.2: third panel). In contrast, few N protein was detected in the BA.1-infected lungs (**Fig. 3B**, second top panel). On day 5 p.i., B.1.1- and BA.5 N proteins were also distributed in the peripheral alveolar space and the higher amount was observed in the group of B.1.1 (**Fig. 3B**). The N proteins were hardly detected in the lungs infected with BA.1 and BA.2. These data suggest that BA.5 efficiently infect bronchial/bronchiolar epithelium and invade the alveolar space more than BA.1 and BA.2. In contrast, BA.1 and BA.2 infect only a portion of bronchial/bronchiolar epithelium and is less efficiently transmitted to the neighboring epithelial cells. Overall, IHC data suggest that BA.5 has a higher spread of infection from the bronchi/bronchioles to the peripheral part of the alveoli than BA.1 and BA.2 but not reaches to the levels of B.1.1.

**Fig. 3.**
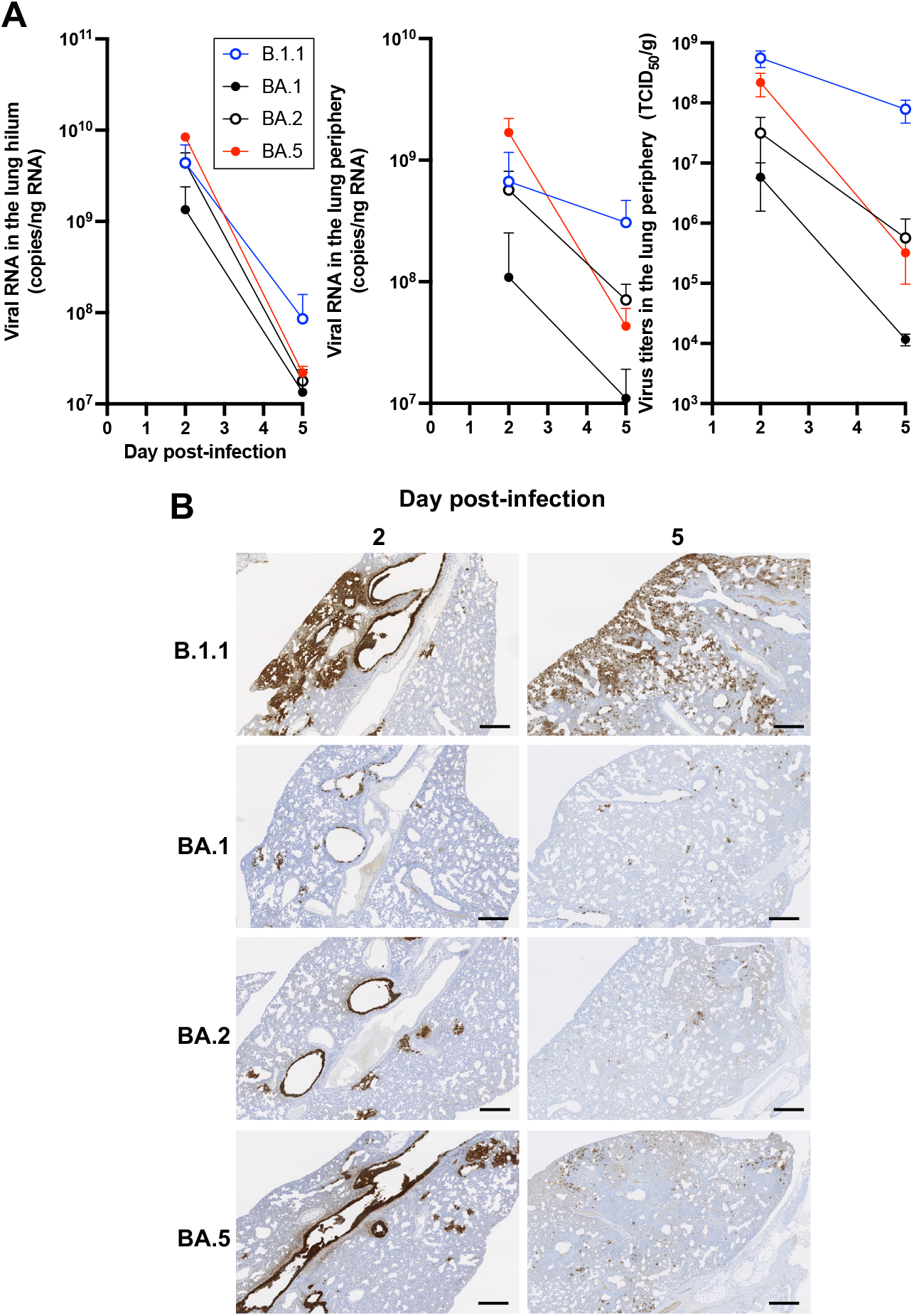
Virological features of Omicron subvariants *in vivo*. Syrian hamsters were intranasally inoculated with B.1.1 (*n* = 4), BA.1 (*n* = 4), BA.2 (*n* = 4), and BA.5 (*n* = 4). (**A**) Viral RNA quantification and titration. Viral RNA load (left, middle) in the lung hilum and periphery and viral titer (right) in the lung periphery were quantified. Statically differences between B.1.1 and other variants were tested with Tukey’s multiplicity correction (‡, *P* < 0.05) and differences between B.1.1 and other variants (*, *P* < 0.05). (**B**) IHC of the SARS-CoV-2 N protein in the lungs of infected hamsters. Representative IHC panels of the viral N proteins in the lower lobe of lungs of the infected hamsters. Scale bars, 500 μm.

## Inflammation in lung tissue infected with Omicron subvariants

To further investigate the pathogenicity of Omicron subvariants in the lung, the formalin-fixed right lungs of infected hamsters were analyzed by carefully identifying the four lobules and main bronchus and lobar bronchi sectioning each lobe along with the bronchial branches. Histopathological scoring was performed as described in the previous study^9–11^. Briefly, pathological features including bronchitis or bronchiolitis, hemorrhage or congestion, alveolar damage with epithelial apoptosis and macrophage infiltration, and hyperplasia of type II pneumocytes were evaluated by certified pathologists and the degree of these pathological findings were arbitrarily scored using four-tiered system as 0 (negative), 1 (weak), 2 (moderate), and 3 (severe). Bronchitis is an inflammatory indicator at early stage of infection. On day 2 p.i., the B.1.1-infected hamsters showed most severe bronchiolitis followed by BA.5-, BA.2-, and BA.1-infected hamsters (**Fig. 4A, B**). On day 5 p.i., the B.1.1-infected hamsters exhibited more severe alveolar damage, hemorrhage, and type II pneumocytes than the animals infected with Omicron variants (**Fig. 4A, 4B**). Between BA.5 and the parental BA.2, infection with BA.5 exhibited more severe inflammation including the alveolar damage as infiltration of lymphocytes and macrophages (score 2.0) and the presence of type II pneumocytes with enlarged cellular cytoplasm and nucleus (score 2.0) compared with BA.2 (both scores 1.75). Higher degree of hemorrhage was observed in BA.5 (2.5) than BA.2 (2.25) (**Fig. 4A**). In case of BA.1-infection, all histological scores were lowest on day 5 p.i. (**Fig. 4A**). Although B.1.1 infection caused most severe inflammation, these data suggest that BA.5 inflammation caused severe inflammation among Omicron subvariants.

**Fig. 4.**
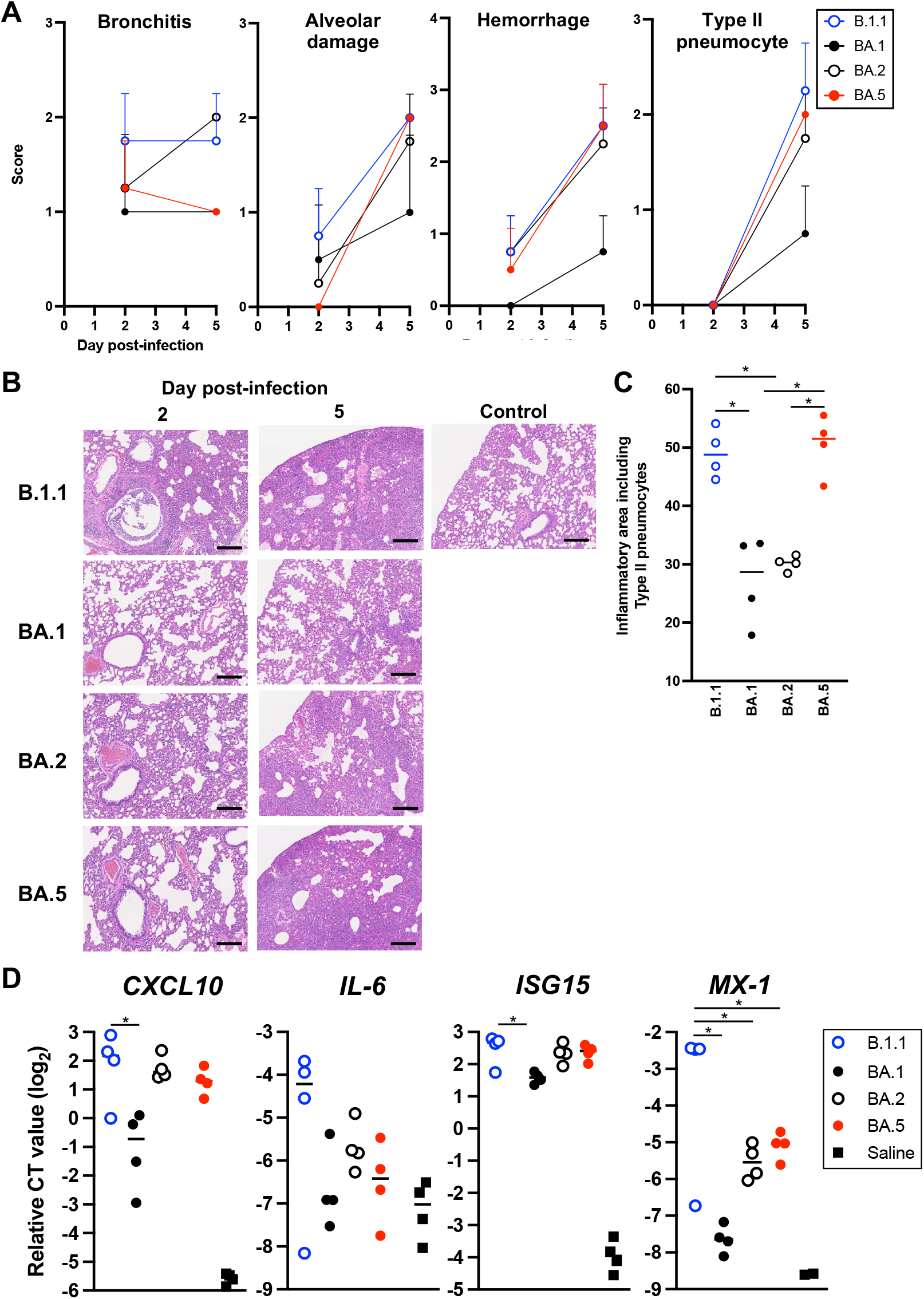
Pathological features of Omicron subvariants. Syrian hamsters were intranasally inoculated with B.1.1 (*n* = 4), BA.1 (*n* = 4), BA.2 (*n* = 4) and BA.5 (*n* = 4). (**A**) Histopathological scoring of lung lesions. (**B**) H&E staining of the lungs of infected hamsters. Uninfected lung alveolar space and bronchioles are also shown. (**C**) Summary of the percentage of the section represented by the inflammatory area with type II pneumocytes. The raw data are shown in **Figure S4**. (**D**) mRNA of the lung tissues obtained on day 2 post infection were used to measure expression levels of inflammatory genes (*CXCL10*, *IL-6*, *ISG15*, and *MX-1*) with normalization of a house-keeping gene, RPL18. In **C**, data are the average ± S.E.M. In **A**, **C**, and **D**, each dot indicates the result from an individual hamster. In **C**, statistically significant differences (*, *P* < 0.05) were determined by two-sided unpaired Student’s *t*-tests without adjustment for multiple comparisons. In **D**, differences among Omicron subvariants were tested with Tukey’s multiplicity correction (*, *P* < 0.05).

Because the difference of histological scoring among the Omicron subvariants were little, inflammatory area mainly composed of the type II pneumocytes with various inflammatory cells as neutrophils, lymphocytes, and macrophages (termed the area of type II pneumocytes) were morphometrically analyzed and found that the area of type II pneumocytes was significantly higher in BA.5 (50.5%) than BA.1 (27.2%) or BA.2 (30.2%) (**Fig. 4C** and **Supplementary Fig. 4**), suggesting that deterioration of inflammation might determine increased pathogenicity of BA.5 in hamsters. Thus, we evaluated inflammatory response upon infection with Omicron subvariants *in vivo,* the mRNA of the lung hilum area on day 2 p.i. and 4 parameters (*CXCL10*, *IL-6*, *ISG15*, and *MX-1*) were measured (**Fig. 4D**). Upon infection with all variants, the evaluated ISGs, *CXCL10*, *ISG15*, and *MX-1*, were upregulated and the expression levels by B.1.1 were highest then followed by the BA.5. The expression of IL-6 was also upregulated and was remained only of B.1.1-infected hamsters but not of Omicron subvariants-infected ones, suggesting that anti-viral inflammatory response led to deterioration of pathogenicity.

## Discussion

Because a new subvariants BA.5 is surged dramatically and outcompeted to the parental BA.1 and BA.2 variants, researchers rapidly dispatched reports showing further resistance of the SARS-CoV-2 Omicron variant against the immunity elicited by previous infections and vaccination^18–23^. In addition, the morbidity of COVID-19 by infection with BA.5 is escalated^3^, suggesting that the appropriate implementation of control measurement is urgently required. However, comparative analysis of these Omicron subvariants, BA.1, BA.2, and BA.5 have not been well documented. Kawaoka et al. showed that the clinical isolate BA.5 exhibited lower pathogenicity than the ancestral Delta in hamster models^24^. We recently reported that the spike protein of BA.5 contributes the enhanced pathogenicity compared to the previous Omicron subvariant BA.2 in our hamster model. Here, we further investigate the *in vitro* and *in vivo* characters of three clinical isolates of Omicron subvariants BA.1, BA.2, and BA.5. Although the virulence of the Omicron subvariants is less than that of the ancestral lineage B.1.1, our comprehensive analyses suggest that the BA.5 gains pathogenicity by evolving to enhance inflammatory response. This might be a key factor for deterioration of mobility in human population by BA.5 infection^3^.

As we showed the series of studies using the recombinant viruses^9–12^, fusogenicity by the viral spike protein has a great impact on viral replication and pathogenesis. Consistent with these, the fusogenicity of the Omicron subvariants were less than the conventional B.1.1 strain and BA.5 exhibited slight high fusogenicity among Omicron subvariants (**Figs. 1C, 1D and 1E**). In addition, *in vitro* growth kinetics is similar in cell lines, VeroE6/TMPRSS2, Calu-3 cells and in iPS cell-derived lung epithelial cells (**Figs. 1A and 1B**). Interestingly however, investigation with airway-on-a-chip as much more mimic *in vivo* environment were conducted and the resulting data showed that BA.5 has a strong barrier disruption capacity among the Omicron subvariants (**Fig. 1F**). In the hamsters, infection with BA.5 showed the strong hemorrhage and alveolar damage (**Figs. 4A and 4B**), supporting the enhanced phenotype of BA.5 for invasion of respiratory tissues.

Kawaoka et al. recently showed that the Omicron subvariants including BA.5 is less pathogenic than Delta and deterioration of weight loss by BA.5 infection is slightly higher than that by BA.2 infection. In the present study, the weight loss in BA.5 infected hamsters was higher than that of other Omicron subvariants, consisting with this. In addition, by using the recombinant virus replacing spike protein gene with the BA.2 backbone, the recombinant virus bearing BA.4/5 spike protein exhibited significantly enhanced pathogenicity in hamsters in our previous study. In combination with our present and previous studies, the possible evolution of BA.5 spike protein to achieve higher pathogenicity shall need to be underscored.

Of these Omicron subvariants, BA.5 exhibited deterioration of weight loss and of respiratory markers (**Fig. 2**). BA.5 isolate efficiently infect bronchial/bronchiolar epithelium and invade the alveolar space leading to the remaining virus replication in the lung. As shown in a clinical report, BA.5 exhibits higher morbidity than BA.2, suggesting inflammation influences outcoming clinical manifestations. Thus, we evaluated the inflammatory responses evoked by viral infection. By morphometrical analysis which is sensitive to reflect the subtle difference of inflammations, the inflammatory area of the lungs infected with BA.5 was larger than that of BA.2 and approximately equal to B.1.1. on day 5 p.i. (**Fig. 4C**), indicating that BA.5 is more immunopathogenic than BA.2. In mRNA levels, interferon-stimulating genes including *CXCL10*, *ISG15*, and *MX-1* were upregulated upon infection (**Fig. 4D**). Reuschl et al., showed the enhanced innate immune suppression by BA.4 and BA.5 compared with the previous BA.2 subvariants^25^. Altogether, severe inflammations of BA.5 might reflect severe clinical manifestations currently happening in human population^3^.

In summary, our analyses using clinically isolated Omicron subvariants, BA.1, BA.2, and BA.5 by *in vitro* experiments and the established animal models contributed to improved understanding of SARS-CoV-2 ecology and evolution. Our findings suggest that the characterization described herein should be offered as an aid in implementing control measurements on COVID-19.

## Methods

### Ethics statement

All experiments with hamsters were performed in accordance with the Science Council of Japan’s Guidelines for the Proper Conduct of Animal Experiments. The protocols were approved by the Institutional Animal Care and Use Committee of National University Corporation Hokkaido University (approval numbers 20-0123 and 20-0060).

### Cell culture

HEK293-ACE2/TMPRSS2 cells [HEK293 cells (ATCC CRL-1573) stably expressing human ACE2 and TMPRSS2]^12^ were maintained in Dulbecco’s modified Eagle’s medium (DMEM) (high glucose) (Nacalai Tesque, Cat# 08459-64) containing 10% fetal bovine serum (FBS) and 1% penicillin-streptomycin (PS). VeroE6/TMPRSS2 cells (VeroE6 cells stably expressing human TMPRSS2; JCRB1819)^15^ were maintained in DMEM (low glucose) (Sigma-aldrich, Cat# D6046-500ML) containing 10% FBS, G418 (1 mg/ml; Nacalai Tesque, Cat# G8168-10ML) and 1% PS. Calu-3 cells (a human airway epithelial cell; ATCC HTB-55) were maintained in EMEM (Wako, Cat# 055-08975) containing 10% FBS and 1% PS. Human alveolar epithelial cells derived from human induced pluripotent stem cells (iPSCs) were manufactured according to established protocols as described below (see “Preparation of human alveolar epithelial cells from human iPSCs” section) and provided by HiLung Inc.

### Preparation of human alveolar epithelial cells from human iPSCs

The air-liquid interface culture of alveolar epithelial cells were differentiated from human iPSC-derived lung progenitor cells as previously described ^26–28^. Briefly, lung progenitor cells were stepwise induced from human iPSCs referring a 21-days and 4-steps protocol ^26^. On day 21, lung progenitor cells were isolated with specific surface antigen carboxypeptidase M and seeded onto upper chamber of 24-well Cell Culture Insert (Falcon, #353104), followed by 28-day and 7-day differentiation of alveolar epithelial cells, respectively. Alveolar differentiation medium supplemented with dexamethasone (Sigma-Aldrich, Cat# D4902), KGF (PeproTech, Cat# 100-19), 8-Br-cAMP (Biolog, Cat# B007), 3-Isobutyl 1-methylxanthine (IBMX), CHIR99021 (Axon Medchem, Cat# 1386), and SB431542 (FUJIFILM Wako, Cat# 198-16543) was used for induction of alveolar epithelial cells. PneumaCult ALI (STEMCELL Technologies, Cat# ST-05001) supplemented with heparin and Y-27632 (LC Laboratories, Cat# Y-5301) hydrocortisone (Sigma-Aldrich, Cat# H0135) was used for induction of airway epithelial cells.

### SARS-CoV-2 preparation and titration

An Omicron subvariants (BA.1 lineage, strain TY38-873, GISAID ID: EPI_ISL_7418017; BA.2 lineage, strain TY40-385, GISAID ID: EPI_ISL_9595859; BA.5 lineage, strain TKYS14631, GISAID ID: EPI_ISL_12812500)^14^ was obtained from the National Institute of Infectious Diseases (BA.1 and BA.2) and Tokyo Metropolitan Institute of Public Health, Japan. An early pandemic D614G-bearing isolate (B.1.1 lineage, strain TKYE610670; GISAID ID: EPI_ISL_479681) was used in the previous study^10^. Virus preparation and titration was performed as previously described^10,12,29^. To prepare the working virus stock, 20 μl of the seed virus was inoculated into VeroE6/TMPRSS2 cells (5 × 10^6^ cells in a T-75 flask). One hour after infection, the culture medium was replaced with DMEM (low glucose) (Wako, Cat# 041-29775) containing 2% FBS and 1% PS. At 3 d.p.i., the culture medium was harvested and centrifuged, and the supernatants were collected as the working virus stock. The viral genome sequences of working viruses were verified as described below.

The titer of the prepared working virus was measured as the 50% tissue culture infectious dose (TCID_50_). Briefly, one day before infection, VeroE6/TMPRSS2 cells (10,000 cells) were seeded into a 96-well plate. Serially diluted virus stocks were inoculated into the cells and incubated at 37°C for 4 d. The cells were observed under microscopy to judge the CPE appearance. The value of TCID_50_/ml was calculated with the Reed–Muench method^30^.

### SARS-CoV-2 infection

One day before infection, VeroE6/TMPRSS2 cells and Calu-3 cells were seeded into a 12-well plate. SARS-CoV-2 was inoculated as a m.o.i. = 0.1 and incubated at 37°C for 1 h. The infected cells were washed, and 1 ml of culture medium was added. The culture supernatant (100 μl) and cells were collected at the indicated time points and purified RNA for RT–qPCR to quantify the viral RNA copy number (see below). To monitor the syncytium formation in infected cell culture, bright-field photos were obtained using an Elipse Ts2 microscope (Nikon).

The infection experiment human iPSC-derived alveolar epithelial cells was performed as previously described^10^. Briefly, the working viruses were diluted with Opti-MEM (Thermo Fisher Scientific, Cat# 11058021). The diluted viruses (1,000 TCID_50_ in 100 μl) were inoculated onto the apical side of the culture and incubated at 37°C for 1 h. The inoculated viruses were removed and washed twice with Opti-MEM. To harvest the viruses on the apical side of the culture, 100 μl Opti-MEM was applied onto the apical side of the culture and incubated at 37°C for 10 min. The Opti-MEM applied was harvested and used for RT–qPCR to quantify the viral RNA copy number (see below).

### Syncytia formation assay

A equal amount of VeroE6/TMPRSS2 cells expressing either EGFP or mCheery ware seeded into the 12-well plate once day before SARS-CoV-2 infection. The infection was conducted as described above and captured the image at 48 h p.i. Florescent images were acquired with Eclipse Ti2 (Nikon, Tokyo, Japan) microscope, equipped with a PlanApo 20×/0.8 objective lens, a TI2-CTRE microscope controller (Nikon), a TI2-S-SE-E motorized stage (Nikon), and an X-Cite turbo system (Excelitas Technologies). The detectors used in this study was a PRIME95B scientific complementary metal-oxide semiconductor (sCMOS) camera (Oxford Instruments). The sets of excitation and emission filters and dichroic mirrors adopted for this observation included GFP HQ (Nikon) for EGFP, Cy3 HQ (Nikon) for mCherry.

### Immunoblotting

Cells lysed on ice in lysis buffer (20 mM Tris-HCl [pH 7.4], 135 mM NaCl, 1% Triton-X 100, 10% glycerol) supplemented with a protease inhibitor cocktail, cOmplete mini (MilliporeSigma), were boiled in loading buffer and subjected to 5%–20% gradient SDS-PAGE. The proteins were transferred to polyvinylidene difluoride membranes (MilliporeSigma) and incubated with the anti-SARAS-CoV-2 S antibody (1:5,000 dilution; GeneTex) or anti-N antibody (1:5,000 dilution; Sino Biological). The immune complexes were visualized with SuperSignal West Femto substrate (Thermo Fisher Scientific). The signals were detected by use of a WSE-LuminoGraph I (ATTO) and ImageSaver6 (ATTO).

### Airway-on-a-chip

Human lung microvascular endothelial cells (HMVEC-L) were obtained from Lonza and cultured with the EGM-2-MV medium (Lonza). To prepare the airway-on-a-chip, first, the bottom channel of a polydimethylsiloxane (PDMS) device was pre-coated with fibronectin (3 μg/ml, Sigma). The microfluidic device was generated according to our previous report^31^. HMVEC-L were suspended at 5×10^6^ cells/mL in the EGM2-MV medium. Then, 10 μl suspension medium was injected into the fibronectin-coated bottom channel of the PDMS device. Then, the PDMS device was turned upside down and incubated for 1 h. After 1 h, the device was turned over, and EGM2-MV medium was added into the bottom channel. After 4 days, human airway organoids (AO) were dissociated and seeded into the top channel. The AO was generated according to our previous report^32^. AO were dissociated into single cells and then suspended at 5×10^6^ cells/mL in AO differentiation medium. Ten μL suspension medium was injected into the top channel. After 1 h, AO differentiation medium was added to the top channel. The cells were cultured under a humidified atmosphere with 5% CO_2_ at 37 °C.

### Electron Microscopy

EM analyses were performed as previously described^33^. Briefly, cells were fixed with 2% glutaraldehyde (Electron Microscopy Sciences) in HEPES buffer (pH 7.4; 30 mM HEPES, 0.1 M NaCl and 2 mM CaCl_2_) and 2% osmium tetroxide (TAAB) in 0.1 M imidazole (pH 7.4). After dehydration and substitution, the cells were embedded in Araldite-Epon (Electron Microscopy Sciences). Ultrathin sections (~70 nm) were mounted on formvar-coated copper grids (SP 3 slit mesh, Nisshin EM), and the sections were imaged with a transmission electron microscope (JEM-1400, JEOL) operating at 80 kV.

### RT–qPCR

RT–qPCR was performed as previously described^10,12,29^. Briefly, 5 μl of culture supernatant was mixed with 5 μl of 2 × RNA lysis buffer [2% Triton X-100, 50 mM KCl, 100 mM Tris-HCl (pH 7.4), 40% glycerol, 0.8 U/μl recombinant RNase inhibitor (Takara-Bio)] and incubated at room temperature for 10 min. RNase-free water (90 μl) was added, and the diluted sample (2.5 μl) was used as the template for real-time RT-PCR performed according to the manufacturer’s protocol using the One Step TB Green PrimeScript PLUS RT-PCR kit (Takara-Bio) and the following primers: Forward *N,* 5’-AGC CTC TTC TCG TTC CTC ATC AC-3’; and Reverse *N*, 5’-CCG CCA TTG CCA GCC ATT C-3’. The viral RNA copy number was standardized with a SARS-CoV-2 direct detection RT-qPCR kit (Takara, Cat# RC300A). Fluorescent signals were acquired using QuantStudio Real-Time PCR system (Thermo Fisher Scientific), CFX Connect Real-Time PCR Detection system (Bio-Rad), Eco Real-Time PCR System (Illumina), qTOWER3 G Real-Time System (Analytik Jena) or 7500 Real-Time PCR System (Thermo Fisher Scientific).

To evaluate Inflammation levels evoked by viral infection in hamsters, 500 ug of the lung RNA were subjected to synthesize cDNA using SuperScript IV VILO Master Mix The resulting cDNA was used to quantify host genes^34^ (see **Supplementary Table 1**) with a Power SYBER Green Master Mix (Thermo Fisher Scientific) and QuantStudio Real-time PCR System (Thermo Fisher Scientific).

### Plasmid construction

The cDNA clones of EGFP and mCherry were inserted between the *XhoI* and *XbaI* sites of the lentiviral vector pCSII-EF-RfA^35^ by using the Infusion technique, and the resulting plasmids were designated pCSII-EF-EGFP, pCSII-EF-mCherry respectively.

### Animal experiments

Syrian hamsters (male, 4 weeks old) were purchased from Japan SLC Inc. (Shizuoka, Japan). Baseline body weights, respiratory parameters, and SpO_2_ were measured before infection. For the virus infection experiments, hamsters were euthanized by intramuscular injection of a mixture of 0.15 mg/kg medetomidine hydrochloride (Domitor^®^, Nippon Zenyaku Kogyo), 2.0 mg/kg midazolam (Dormicum^®^, FUJIFILM Wako Chemicals) and 2.5 mg/kg butorphanol (Vetorphale^®^, Meiji Seika Pharma) or 0.15 mg/kg medetomidine hydrochloride, 2.0 mg/kg alphaxaone (Alfaxan^®^, Jurox) and 2.5 mg/kg butorphanol. The B.1.1 virus, Omicron subvariants (5 × 1,000 TCID_50_ in 100 μl), or saline (100 μl) were intranasally inoculated under anesthesia. Oral swabs were collected at indicated timepoints. Body weight was recorded daily by day 7 p.i., and days 10 and 14 p.i. Enhanced pause (Penh, see below), the ratio of time to peak expiratory follow relative to the total expiratory time (Rpef, see below), and subcutaneous oxygen saturation (SpO_2_, see below) were monitored on days 1, 3, 5, and 7 p.i. Lung tissues were anatomically collected on days 2 and 5 .p.i. Viral RNA load in the oral swabs and respiratory tissues were determined by RT–qPCR. Viral titers in the lung periphery were determined by a TCID_50_. These tissues were also used for histopathological and IHC analyses (see below).

### Lung function test

Respiratory parameters (Penh and Rpef) were measured by using a whole-body plethysmography system (DSI) according to the manufacturer’s instructions. In brief, a hamster was placed in an unrestrained plethysmography chamber and allowed to acclimatize for 30 s, then, data were acquired over a 2.5-m period by using FinePointe Station and Review softwares v2.9.2.12849 (STARR). The state of oxygenation was examined by measuring SpO_2_ using pulse oximeter, MouseOx PLUS (STARR). SpO_2_ was measured by attaching a measuring chip to the neck of hamsters sedated by 0.25 mg/kg medetomidine hydrochloride.

### H&E staining

H&E staining was performed as described in the previous study^10^. Briefly, excised animal tissues were fixed with 10% formalin neutral buffer solution and processed for paraffin embedding. The paraffin blocks were sectioned with 3 μm-thickness and then mounted on silane-coated glass slides (MAS-GP, Matsunami). H&E staining was performed according to a standard protocol.

### IHC

IHC was performed using an Autostainer Link 48 (Dako). The deparaffinized sections were exposed to EnVision FLEX target retrieval solution high pH (Agilent, Cat# K8004) for 20 m at 97°C to activate, and mouse anti-SARS-CoV-2 N monoclonal antibody (R&D systems, Clone 1035111, Cat# MAB10474-SP, 1:400) was used as a primary antibody. The sections were sensitized using EnVision FLEX (Agilent) for 15 m and visualized by peroxidase-based enzymatic reaction with 3,3’-diaminobenzidine tetrahydrochloride as substrate for 5 m.

For the evaluation of the N protein positivity in the tracheae on day 2 p.i. and the lung specimens of infected hamsters on days 2 and 5 p.i. (B1.1, BA.1, BA.2, and BA.5, n = 4 each) were stained with mouse anti-SARS-CoV-2 N monoclonal antibody (R&D systems, Clone 1035111, Cat# MAB10474-SP, 1:400). The N protein positivity was evaluated by certificated pathologists as previously described ^9,11^. Images were incorporated as virtual slide by NDP.scan software v3.2.4 (Hamamatsu Photonics). The N-protein positivity was measured as the area using Fiji software v2.2.0 (ImageJ).

### Histopathological scoring of lung lesion

The inflammation area in the infected lungs was measured by the presence of the type II pneumocyte hyperplasia. Four hamsters infected with each virus were sacrificed on days 2 and 5 p.i., and all four lung lobes, including right upper (anterior/cranial), middle, lower (posterior/caudal), and accessory lobes, were sectioned along with their bronchi. The tissue sections were stained by H&E, and the digital microscopic images were incorporated into virtual slides using NDRscan3.2 software (Hamamatsu Photonics). The color of the images was decomposed by RGB in split channels using Fiji software v2.2.0.

Histopathological scoring was performed as described in the previous study. Briefly, pathological features including bronchitis or bronchiolitis, hemorrhage or congestion, alveolar damage with epithelial apoptosis and macrophage infiltration, hyperplasia of type II pneumocytes, and the area of the hyperplasia of large type II pneumocytes were evaluated by certified pathologists and the degree of these pathological findings were arbitrarily scored using four-tiered system as 0 (negative), 1 (weak), 2 (moderate), and 3 (severe). The “large type II pneumocytes” are the hyperplasia of type II pneumocytes exhibiting more than 10-μm-diameter nucleus. Total histology score is the sum of these five indices. In the representative lobe of each lung, the inflammation area with type II pneumocytes was gated by the certificated pathologists on H&E staining, and the indicated area were measured by Fiji software v2.2.0.

### Viral genome sequencing analysis

The sequences of the working viruses were verified by viral RNA-sequencing analysis. Viral RNA was extracted using QIAamp viral RNA mini kit (Qiagen, Cat# 52906). The sequencing library for total RNA-sequencing was prepared using NEB Next Ultra RNA Library Prep Kit for Illumina (New England Biolabs, Cat# E7530). Paired-end, 76-bp sequencing was performed using MiSeq (Illumina) with MiSeq reagent kit v3 (Illumina, Cat# MS-102-3001). Sequencing reads were trimmed using fastp v0.21.0^36^ and subsequently mapped to the viral genome sequences of a lineage B isolate (strain Wuhan-Hu-1; GISAID ID: EPI_ISL_402125; GenBank accession no. NC_045512.2) using BWA-MEM v0.7.17^37^. Variant calling, filtering, and annotation were performed using SAMtools v1.9^38^ and snpEff v5.0e^39^. Information on the detected mutations in the working virus stocks is summarized in **Supplementary Table 2**.

### Statistics

Viral RNA copy, body wight, Penh, Rpef, and SpO_2_, and inflammatory mRNA gene levels obtained from the in vivo experiments were analyzed by repeated measures analysis of variance. Inflammation measures upon infection in vivo, the mRNA of the lung hilum area on day 2 p.i. and 4 parameters (CXCL10, IL-6, ISG15, and Mx-1) were compared among Omicron subvariants using analysis of variance. Regarding Penh, Rpef and SpO_2_, we compared between infected and uninfected animals and calculated p-value using Dunnett adjustment. The other measurements were tested with Tukey’s multiplicity correction to maintain type-I error rate to compare among infected or uninfected animals. These analyses were conducted using SAS Ver 9.4 (Cary, NC). Two-sided significance level was set to 0.05. Statistical differences between BA.5 and other variants or saline across timepoints from day 1 p.i. to day 7 p.i. were determined by multiple regression. The family-wise error rates calculated using the Holm method. The indicated analyses were performed in R v4.1.2 (https://www.r-project.org/).

---

**Supplementary Table 1. Primers used for quantifying the host genes of hamster upon infection with the SARS-CoV-2 variants in this study.**

**Supplementary Table 2. Mutations in Omicron subvariants.**

**Supplementary Fig. 1.**
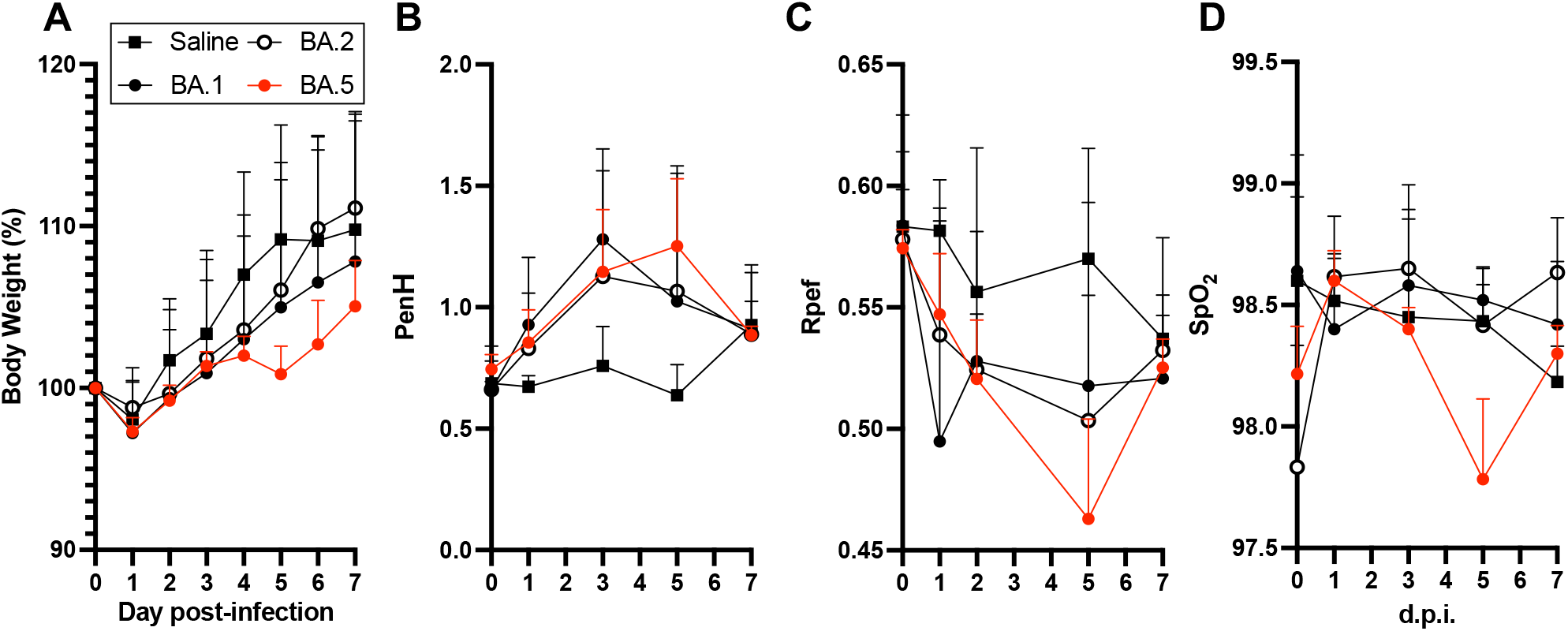
*In vivo* dynamics of Omicron subvariants in related to Fig.2. Dynamics of of body weight (A), Penh (B), Rpef (C), and SpO2 (D) within Omicron subvariants by days 7 p.i were shown. Saline injection was served as a control in this study.

**Supplementary Fig. 2.**
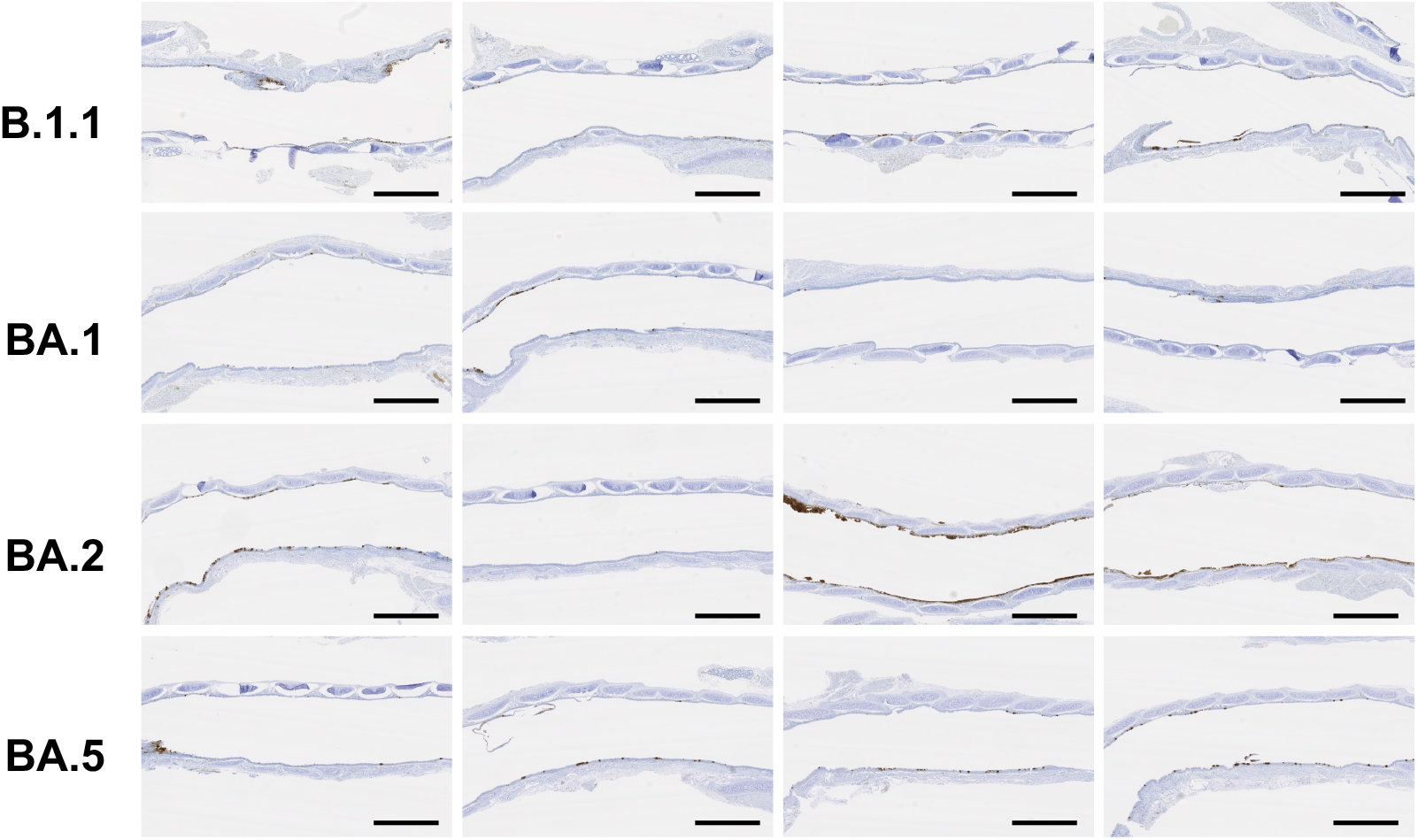
Virological features of Omicron subvariants *in vivo* in the trachea related to Fig. 3. Percentage of N-positive cells in the upper part of the trachea from the oral entrance at the vertical levels of thyroid cartilage on day 2 p.i. was shown. Scale bars, 1 mm.

**Supplementary Fig. 3.**
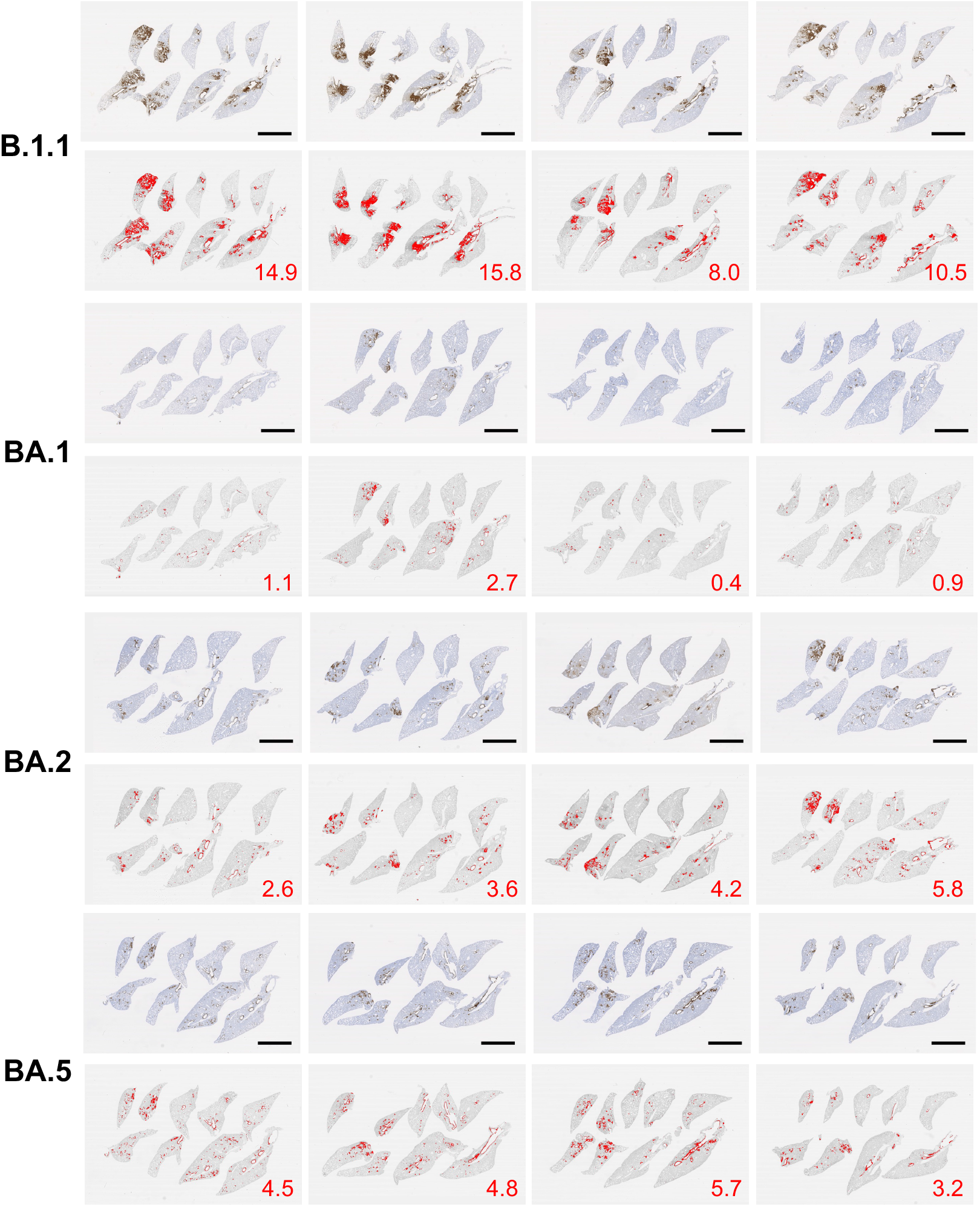
Virological features of Omicron subvariants *in vivo* in the lung related to Fig. 3. Percentage of N-positive cells in whole lung lobes on day 2 p.i. was shown. Scale bars, 5 mm.

**Supplementary Fig. 4.**
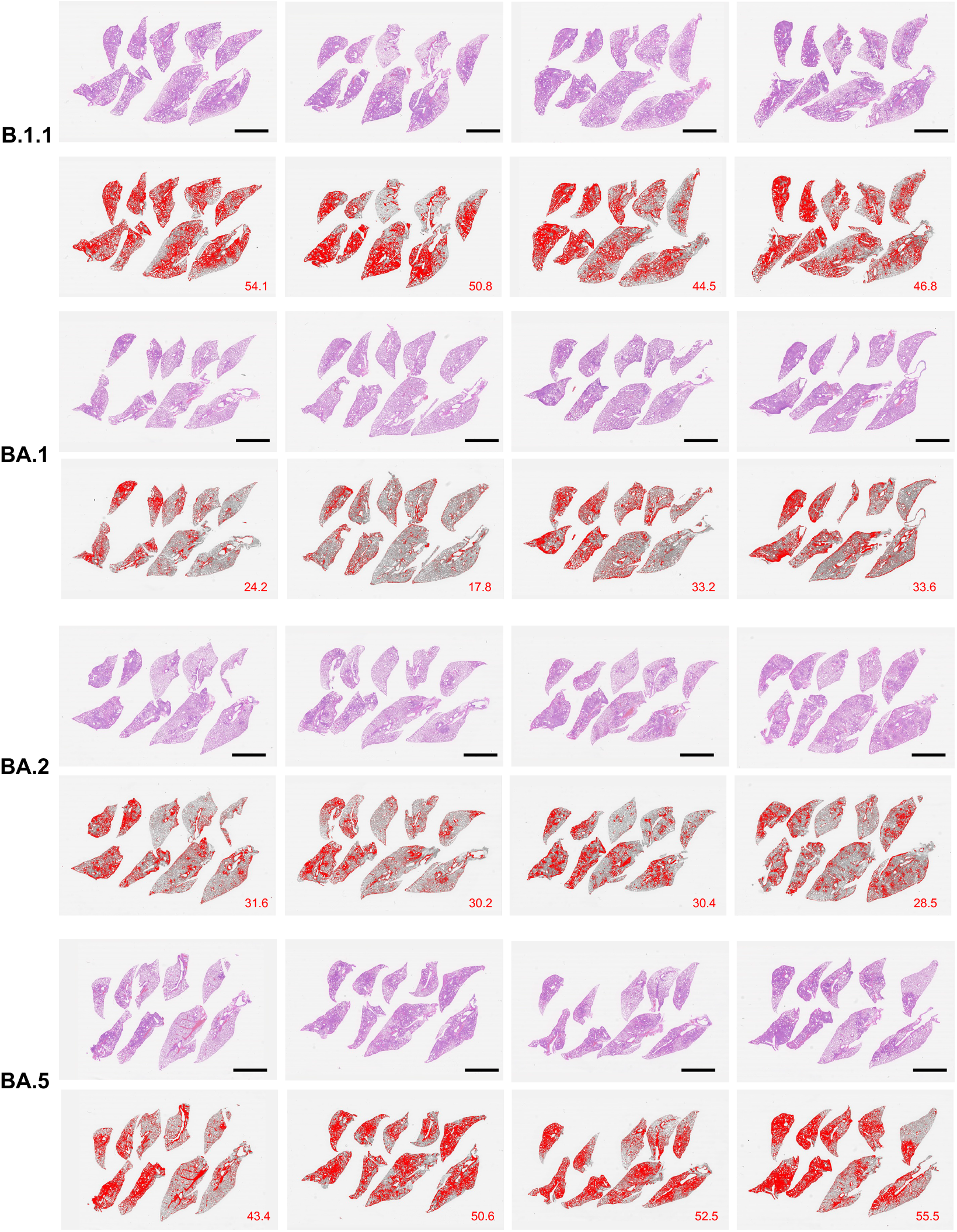
Pathological features of Omicron subvariants *in vivo* related to Fig. 4. Section of all four lung lobes on day 5 p.i. H&E staining and the inflammatory area with type II pneumocytes are shown. The inflammatory area is colored in red. The number in the panel indicates the percentage of the section represented by the indicated area. Scale bars,5 mm.

## Author Contributions

Tomokazu Tamura, Daichi Yamasoba, Tomoko Kamasaki, Rina Hashimoto, Yoichiro Fujioka, Rigel Suzuki, Hayato Ito, Izumi Kimura, performed cell culture experiments.

Tomokazu Tamura, Naganori Nao, Rigel Suzuki, Hayato Ito Mai Kishimoto, Hirofumi Sawa, Kumiko Yoshimatsu performed animal experiments.

Yoshitaka Oda, Lei Wang, Masumi Tsuda, Shinya Tanaka performed histopathological analysis.

Daichi Yamasoba, Izumi Kimura, and Jumpei Ito performed viral genome sequencing analysis.

Jumpei Ito, Isao Yokota performed statistical, modelling, and bioinformatics analyses.

Keita Matsuno, Kazuo Takayama, Kei Sato, Takasuke Fukuhara, designed the experiments and interpreted the results.

Tomokazu Tamura, Takasuke Fukuhara wrote the original manuscript.

All authors reviewed and proofread the manuscript.

The Genotype to Phenotype Japan (G2P-Japan) Consortium contributed to the project administration.

## Acknowledgments

We would like to thank all members belonging to The Genotype to Phenotype Japan (G2P-Japan) Consortium. We thank the National Institute for Infectious Diseases, Japan, for providing BA.1 and BA.2 isolates, and Tokyo Metropolitan Institute of Public Health for providing a BA.5 isolate.

This study was supported in part by AMED Research Program on Emerging and Re-emerging Infectious Diseases (JP20fk0108401, to Takasuke Fukuhara; JP20fk010847, to Takasuke Fukuhara; JP21fk0108617 to Takasuke Fukuhara; JP20fk0108146, to Kei Sato; JP20fk0108451, to G2P-Japan Consortium, Keita Matsuno, Kei Sato, and Takasuke Fukuhara; JP21fk0108494 to G2P-Japan Consortium, Keita Matsuno, Shinya Tanaka, Kei Sato and Takasuke Fukuhara); AMED Program on R&D of new generation vaccine including new modality application (JP223fa727002, to Kei Sato); AMED Research Program on HIV/AIDS (21fk0410039, to Kei Sato); AMED Japan Program for Infectious Diseases Research and Infrastructure (JP22wm0125008, to Hirofumi Sawa and JP21wm0225003, to Hirofumi Sawa); AMED Research Program on infectious disease drug development (22gm1610005h0002, to Kazuo Takayama); JST CREST (JPMJCR20H4, to Kei Sato); JSPS KAKENHI Grant-in-Aid for Scientific Research B (21H02736, to Takasuke Fukuhara); JSPS Core-to-Core Program (A. Advanced Research Networks) (JPJSCCA20190008, to Kei Sato); World-leading Innovative and Smart Education (WISE) Program 1801 from the Ministry of Education, Culture, Sports, Science and Technology (MEXT) (to Naganori Nao); The Tokyo Biochemical Research Foundation (to Kei Sato).

## Data availability

The accession numbers of viral sequences used in this study are listed in Method section.

## Consortia

Marie Kato, Zannatul Ferdous, Hiromi Mouri, Kenji Shishido, Naoko Misawa, Keiya Uriu, Yusuke Kosugi, Shigeru Fujita, Mai Suganami, Mika Chiba, Ryo Yoshimura, So Nakagawa, Jiaqi Wu, Akifumi Takaori-Kondo, Kotaro Shirakawa, Kayoko Nagata, Yasuhiro Kazuma, Ryosuke Nomura, Yoshihito Horisawa, Yusuke Tashiro, Yugo Kawai, Takashi Irie, Ryoko Kawabata, Terumasa Ikeda, Hesham Nasser, Ryo Shimizu, MST Monira Begum, Otowa Takahashi, Kimiko Ichihara, Takamasa Ueno, Chihiro Motozono, Mako Toyoda, Akatsuki Saito, Yuri L. Tanaka, Erika P. Butlertanaka, Maya Shofa, Takao Hashiguchi, Teteki Suzuki, Kanako Kimura, Jiei Sasaki, Yukari Nakajima, Kaori Tabata

